# Two-Photon Imaging of Striatum Demonstrates Distinct Functions for Striosomes and Matrix in Reinforcement Learning

**DOI:** 10.1101/199497

**Authors:** Bernard Bloem, Rafiq Huda, Mriganka Sur, Ann M. Graybiel

## Abstract

Despite the discovery of striosomes several decades ago, technical difficulties have hampered the study of their functions. Here we used 2-photon calcium imaging in neuronal birthdate-labeled Mash1-CreER mice to image simultaneously the activity of striosomal and matrix neurons *in vivo*. We report that with this method we can visually identify circumscribed zones of neuropil that correspond to striosomes as verified in immunostained sections. We find that striosomal neurons, relative to matrix neurons, preferentially encode reward-predicting cues, and that their activity contains more information about expected outcome. These characteristics emerge during training and further strengthen during overtraining. Both striatal compartments are active similarly after reward delivery, firing at neuron-specific times during or after consummatory licking. Finally, we find that immediate reward history strongly modulates neuronal activation in the next trial, especially in matrix neurons. These results suggest that striosomes and matrix have distinct functions in relation to reinforcement learning.

## Introduction

The striatum, despite its relatively homogeneous appearance in simple cell stains, is made up of a mosaic of macroscopic zones, the striosomes and matrix, which differ in their input and output connections and are thought to allow specialized processing by physically modular groupings of striatal neurons (*Crittenden et al., 2016; Fujiyama et al., 2011; Gerfen, 1984; Graybiel and Ragsdale, 1978; Jiménez-Castellanos and Graybiel, 1989; Langer and Graybiel, 1989; Lopez-Huerta et al., 2016; Salinas et al., 2016; Smith et al., 2016; Stephenson-Jones et al., 2016; Walker et al., 1993; Watabe-Uchida et al., 2012*). Particularly striking among these modules are the striosomes (also called patches), which are distinct from the surrounding matrix by their differential expression of neurotransmitters, receptors and many other gene expression patterns (*Banghart et al., 2015; Brimblecombe and Cragg, 2015, 2017; Crittenden and Graybiel, 2011; Cui et al., 2014; Gerfen, 1992; Graybiel, 2010; Graybiel and Ragsdale, 1978*). Striosomes in the anterior striatum have strong inputs from regions related to the limbic system, including parts of the orbitofrontal and medial prefrontal cortex (*Eblen and Graybiel, 1995; Friedman et al., 2015; Gerfen, 1984; Ragsdale and Graybiel, 1990*) and, at subcortical levels, the bed nucleus of the stria terminalis (*Smith et al., 2016*) and basolateral amygdala (*Ragsdale and Graybiel, 1988*). The striosomes are equally specialized in their outputs: they project directly to the dopamine-containing neurons of the substantia nigra (*Crittenden et al., 2016; Fujiyama et al., 2011*) and, via a disynaptic pathway, to the lateral habenula (*Rajakumar et al., 1993; Stephenson-Jones et al., 2016*). By contrast, the matrix and its constituent matrisomes receive abundant input from sensorimotor and associative parts of the neocortex (*Flaherty and Graybiel, 1994; Gerfen, 1984; Parthasarathy et al., 1992; Ragsdale and Graybiel, 1990*), and project into the main direct and indirect pathways to the pallidum and non-dopaminergic pars reticulata of the substantia nigra (*Flaherty and Graybiel, 1994; Giménez-Amaya and Graybiel, 1991; Kreitzer and Malenka, 2008*), universally thought to modulate movement control (*Albin et al., 1989; Alexander and Crutcher, 1990; DeLong, 1990*).

This contrast in connectivity between striosomes and matrix highlights the possibility that striosomes, which physically form three-dimensional labyrinths within the much larger matrix, could serve as limbic outposts within the large sensorimotor matrix. The question of what the actual functions of striosomes are, however, remains unsolved. Answering this question has importance for clinical work as well as for basic science: striosomes have been found, in post-mortem studies, to be selectively vulnerable in disorders with neurologic and neuropsychiatric features (*Crittenden and Graybiel, 2016; Saka et al., 2004; Sato et al., 2008; Tippett et al., 2007*). Ideas about the functions of striosomes have ranged from striosomes serving as the critic in actor-critic architecture models (*Doya, 1999*) to their generating responsibility signals in hierarchical learning models (*Amemori et al., 2011*), to their being critical to motivationally demanding decision-making prior to action (*Friedman et al., 2017; Friedman et al., 2015*), and to other functions (*Brown et al., 1999*). However, the technical difficulties involved in reliably identifying and recording the activity of striosomal neurons, as compared to that of matrix neurons, have been exceedingly challenging; striosomes are too small to yet be detected by fMRI, and their neurons have remained unrecognizable in *in vivo* electrophysiological studies with the exception of those identifying putative striosomes by combinations of antidromic and orthodromic stimulation (*Friedman et al., 2017; Friedman et al., 2015*). With the development of endoscopic calcium imaging (*Bocarsly et al., 2015; Carvalho Poyraz et al., 2016; Luo et al., 2011*) and 2-photon imaging of deep-lying structures (*Dombeck et al., 2010; Howe and Dombeck, 2016; Kaifosh et al., 2013; Lovett-Barron et al., 2014; Mizrahi et al., 2004; Sato et al., 2016*), identifying functions of these specialized striatal zones should be within reach, especially when combined with the use of genetic mouse models that allow direct visual identification of selectively labeled neurons.

Here we report that we have developed a 2-photon microscopy protocol for simultaneously examining the activity of striosomal and matrix neurons in the dorsal caudoputamen of behaving head-fixed mice birthdate-tagged to preferentially label striosomal neurons (*Fishell and van der Kooy, 1987; Graybiel, 1984; Graybiel and Hickey, 1982; Kelly et al., 2017; Newman et al., 2015; Taniguchi et al., 2011*). Key to this work was to achieve dense, permanent labeling of not only striosomal cell bodies, but also their striosome-bounded neuropil. We accomplished this differential labeling by using mice that were pulse-labeled during the generation time of the spiny projection neurons (SPNs) of striosomes. Striosomal neurons were tagged by fate-mapping of neural progenitor cells using Mash(Ascl1)-CreER driver lines with tamoxifen induction at E11.5 (*Kelly et al., 2017*). This method allowed striosomal detection based on the labeling of SPN cell bodies as well as the rich neuropil labeling of the striosomes, capitalizing on the fact that SPN processes of striosome and matrix compartments rarely cross striosomal borders (*Bolam et al., 1988; Lopez-Huerta et al., 2016; Walker et al., 1993*). With the restricted neuropil of the birthdate-labeled cells, it was possible to identify cells as being inside striosomes, and concomitantly to clearly identify neurons outside of the zones of neuropil labeling.

With this method, we compared the activity patterns of striosomal and matrix neurons related to multiple elementary aspects of striatal encoding as mice performed a classical conditioning task. By having cues signaling different reward delivery probabilities, we tested whether striosomes and matrix differentially encode changes in expected outcome and received rewards (*Amemori et al., 2015; Bayer and Glimcher, 2005; Bromberg-Martin and Hikosaka, 2011; Friedman et al., 2015; Keiflin and Janak, 2015; Matsumoto and Hikosaka, 2007; Oyama et al., 2010; Oyama et al., 2015; Schultz, 2016; Schultz et al., 1997; Stalnaker et al., 2012; Watabe-Uchida et al., 2017; Watabe-Uchida et al., 2012*). By imaging day by day during the acquisition and overtraining periods of the task, we asked whether these patterns changed in systematic ways with experience. Finally, we tested the effect of reward history on the activity patterns of current trials, given reports that strong reward-history activity has been found in sites considered to be directly or indirectly connected with striosomes (*Bromberg-Martin et al., 2010; Hamid et al., 2016; Tai et al., 2012*).

We demonstrate that neurons within visually identified striosomes encode reward-predicting tones more strongly than do those of the surrounding matrix, but that matrix neurons are more strongly modulated by reward history, especially during extended post-reward periods. These activity patterns develop during learning with different dynamics for cue responses and post-reward responses. These findings suggest that neurons in striosomes and matrix can be differentially tuned by reinforcement contingencies both during learning and during subsequent performance. Finally, this work opens the opportunity for future functional understanding of striosome-matrix architecture by 2-photon microscopy and selective tagging of neurons with known developmental origins, an opportunity that will be valuable in the study of both normal animals and those representing models of disease states.

## Results

To detect striosomes, we performed experiments in Mash1-CreER X Ai14 mice injected with tamoxifen at E11.5. This pulse labeling method (*Kelly et al., 2017*) resulted in strong tdTomato labeling of clusters of SPNs and surrounding regions of neuropil in the striatum (Figure 1). In initial experiments, we confirmed that these clusters corresponded to striosomes by the close match between the zones of tdTomato neuropil labeling and mu-opioid receptor 1 (MOR1)-rich zones observed in immunohistologically prepared sections (*Tajima and Fukuda, 2013*). We also observed some tdTomato-labeled cells outside of MOR1-labeled striosomes, scattered sparsely in the extra-striosomal matrix.

**Figure 1.**
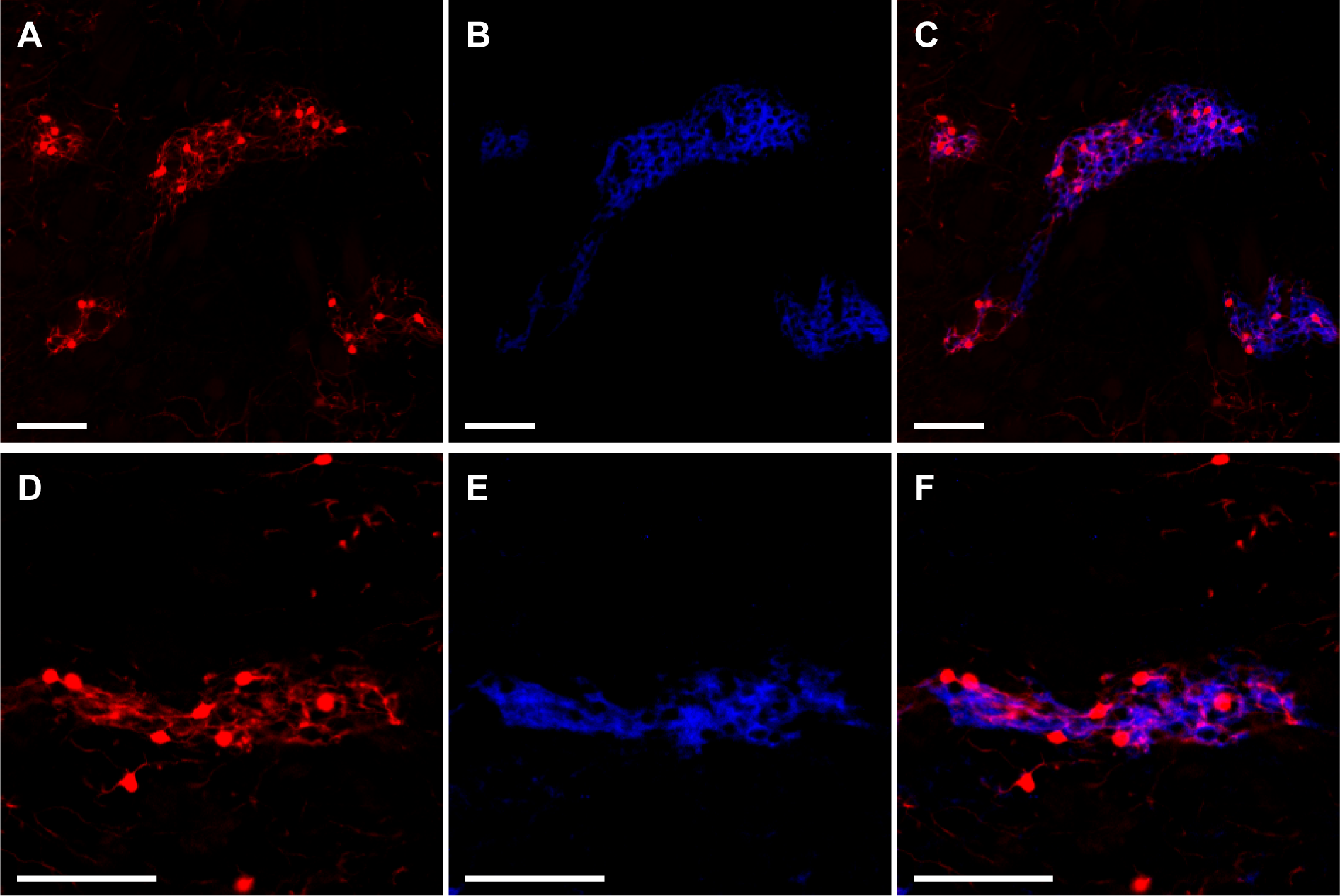
Striosomes are labeled with tdTomato in Mash1-CreER X Ai14 mice that received tamoxifen at E11.5. Images illustrate two examples (rows) of striosomal labeling of cell bodies and neuropil by tdTomato (**A,D,** red) as verified by MOR1 immunostaining (**B,E,** blue) with merges of these marker images (**C,F**). Scale bars represent 100 μm.

For *in-vivo* experiments, we used 2-photon microscopy to image the striatum of 5 striosome-labeled mice that had received unilateral intrastriatal injections of AAV5-hSyn-GCaMP6s and had been implanted with cannula windows and a headplate (Figure 2). Each mouse was trained on a classical conditioning task in which 2 auditory tones (1.5-s duration each) were associated with reward delivery by different probabilities (tone 1, 80% vs tone 2, 20%) (Figure 2A). Inter-trial intervals were 7 ± 1.75 s. With training, mice began to lick in anticipation of the reward, and the amount of this anticipatory licking became greater when cued by the tone indicating a high probability (80%) of reward (Figure 2B). We calculated a learning criterion based on the anticipatory lick rates during the two cues and the subsequent delay period (0.5 s). Mice exhibiting a divergence in anticipatory licking for the two cues for at least two out of three consecutive sessions were considered as trained (Figure 2C). We performed imaging during training (task acquisition) and after this criterion had been reached (trained, Figure 2D).

**Figure 2.**
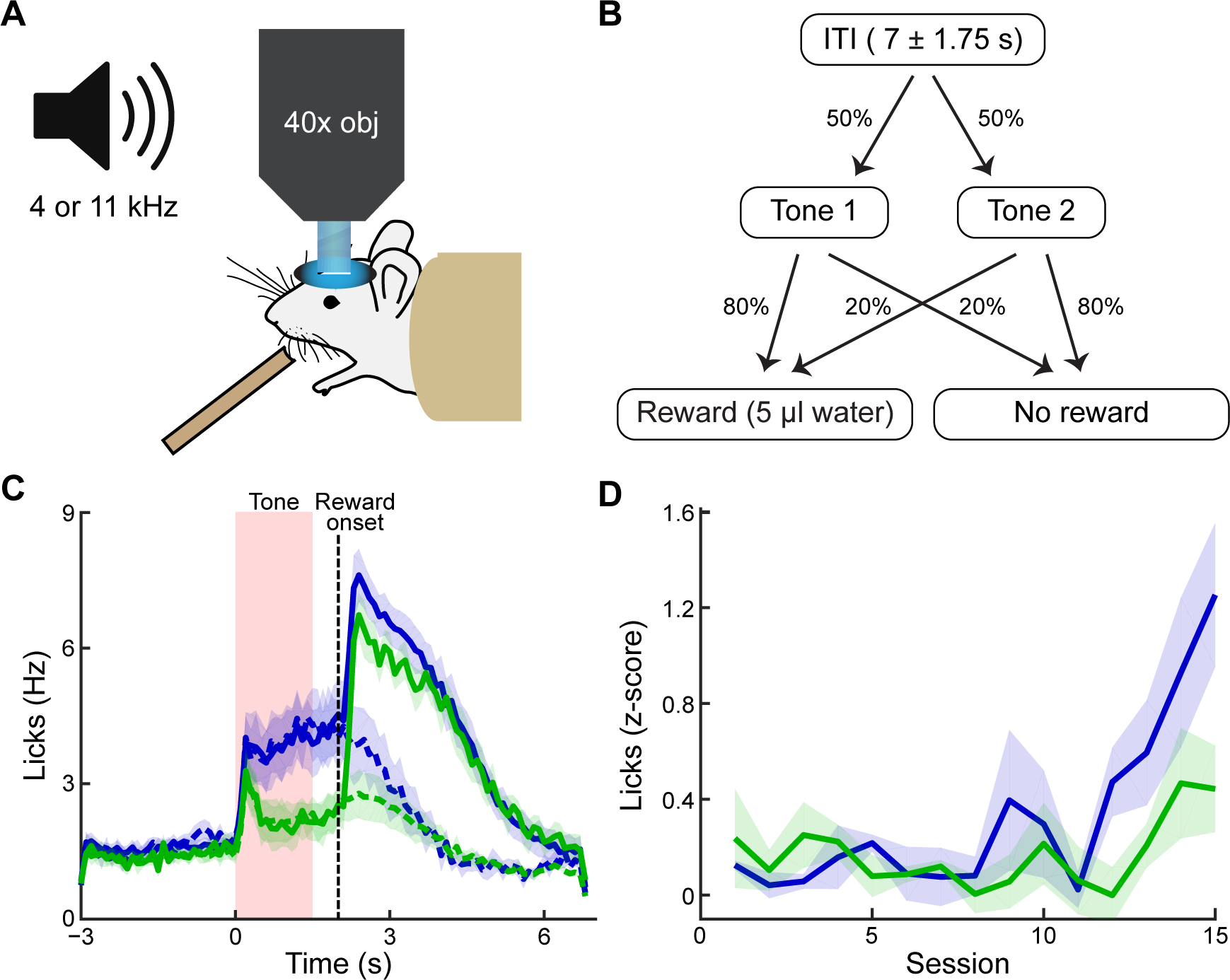
Behavioral task and performance. (**A**) The striatum was imaged during conditioning sessions in which tones predicted reward delivery. (**B**) Two tones (4 and 11 kHz) were played (1.5 s duration) with distinct reward probabilities (80% or 20%). After a 0.5-s delay, reward could be delivered. Inter-trial interval durations varied from 5.25 to 8.75 s. (**C**) Average (± SEM) frequency of licking by trained mice. Anticipatory licking was significantly higher during sounding of the high-probability tone (blue) than during sounding of the low-probability tone (green). After reward delivery, licking rates were elevated for several seconds (solid lines: rewarded trials; dotted lines: unrewarded trials). (**D**) Mice began to exhibit differences in levels of anticipatory licking between the two cues after 11-12 sessions. Animals were considered to be trained when they had 2 out of 3 consecutive sessions with significantly higher anticipatory licking during the high-probability tone (blue). Shading represents SEM.

### Imaging of striosomes

Clusters of tdTomato-positive neurons were clearly visible *in vivo* in the 2-photon microscope at 40x magnification, and the neuropil of these neurons delimited zones in which many dendritic processes could be identified (Figure 3). We simultaneously recorded striosomal and matrix neurons from fields of view with clear striosomes. In all animals, we could see at least two different striosomes from which we imaged at least five different non-overlapping fields of view. In some instances, we could see two different striosomes in one field of view. In the entire data set, we imaged 1867 neurons in striosomes and 4453 in the matrix. Because striosomes form parts of extended branched labyrinths, it was possible to follow some striosomes through ±100 μm in depth, and across ± 800 μm in the field of view. During training, we rotated through the fields of view, but after training criterion had been reached, we recorded activity in unique non-overlapping fields of view (2704 neurons, of which 727 were in striosomes; between 252 -782 neurons per mouse).

**Figure 3.**
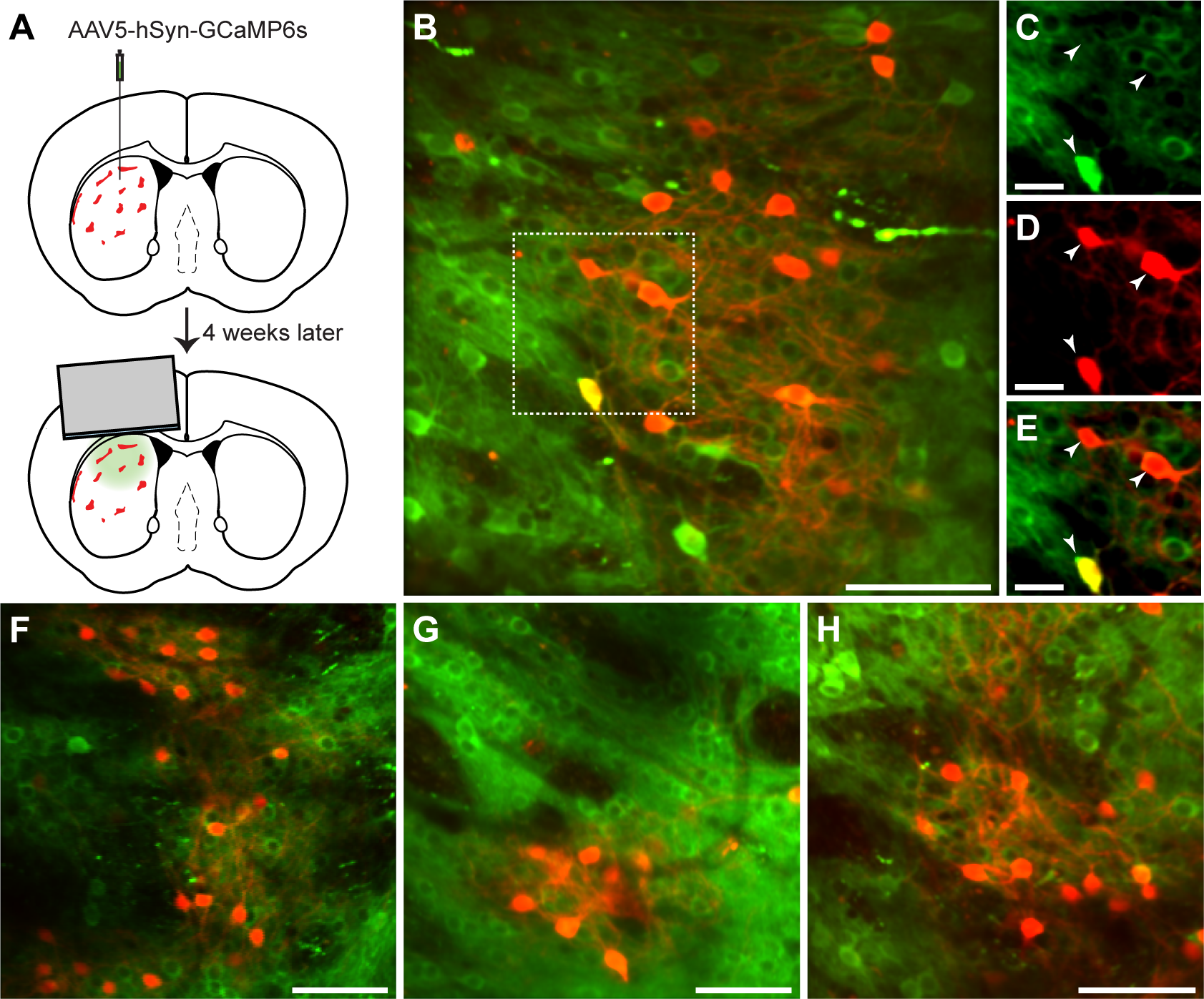
*In vivo* 2-photon calcium imaging of identified striosomes and matrix. (**A**) Mash1-CreEr X Ai14 mice were injected with AAV5-hSyn-GCaMP6s and 4 weeks later were implanted with a cannula. (**B**) Image of a striosome viewed through the 2-photon microscope showing tdTomato labeling in red and GCaMP in green (scale bar: 100 μm) in the striatum of a trained mouse. (**C-E**) Higher magnification of the region indicated in B (scale bar: 10 μm). Arrowheads indicate double-labeled cells. (**F-H**) Representative examples of striosomes imaged in three other trained mice (scale bars: 100 μm).

### Striatal neurons exhibit heightened activity during different task epochs

As an initial approach to our data, we analyzed the overall fluorescence for every session in trained animals by averaging the frame-wide fluorescence (Figure 4A). Both cues evoked a large response in the neuropil signal, which was larger for the high-probability cue. After reward delivery, there was a prolonged, strong activation that peaked around 5 s after reward delivery. To determine more precisely the nature of this activation, we aligned neuronal responses in the rewarded trials to the tone onset, to the first lick after reward delivery and to the end of the licking bout (Figure 4B). This analysis demonstrated that, in addition to the tone response, there was an increase in signal after the start of the consummatory licking period, and this signal increased over time and peaked at the time of the last lick, and then subsided.

**Figure 4.**
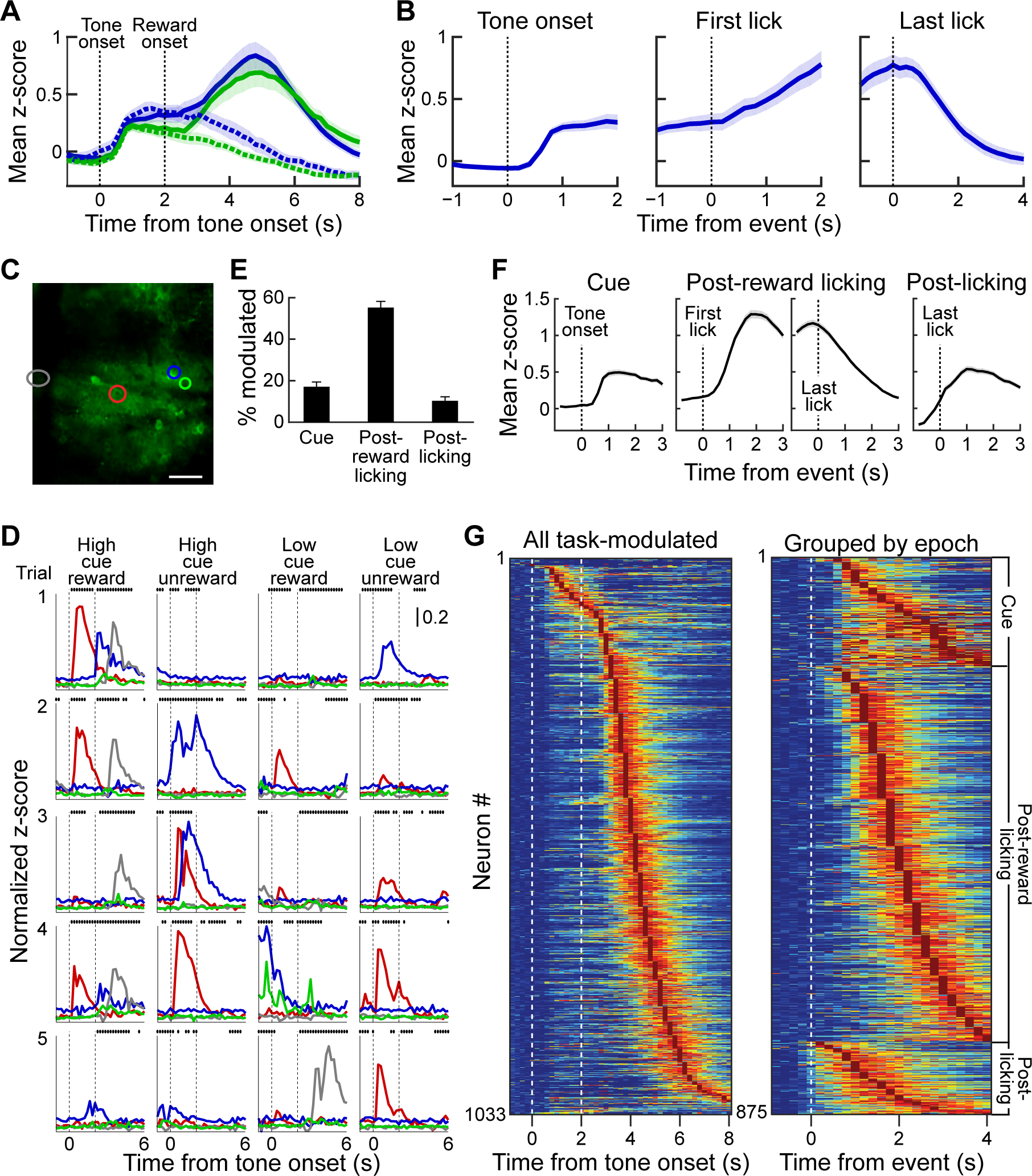
Striatal activity during reward predicting cues and during post-reward period. (**A**) Aggregate neuropil calcium signal in all 4 trial types (blue: high-probability cue; green: low-probability cue; solid line: rewarded trials; dotted line: unrewarded trials). Shading represents SEM. (**B**) Neuropil activation aligned to tone onset (left), first lick after reward delivery (middle) and last lick (right). Only rewarded trials with high-probability cues are included. (**C-D**) Responses of the neurons (**D**) color-coded in C during five sample trials (rows) for four different cue-outcome conditions (columns). Dotted lines indicate the tone and reward onsets. Scale bar in C represents 100 μm. Dots above each plot show when licks occurred. (**E**) Percentage of task-modulated neurons that were selectively active during cue, post-reward licking, or post-licking epochs of the task. Error bars represent 95% confidence intervals. (**F**) Population-averaged responses of task-modulated neurons selectively active during the three epochs. Data for neurons active during the post-reward licking period are aligned to both the first and last lick. (**G**) Session-averaged activity of all task-modulated neurons (left) and those that were significantly active during only one of three task epochs (right). Neurons were sorted by the timing of their peak activity.

Next, we analyzed single-cell activity to investigate the neural dynamics of task encoding by the striatal neurons. In particular, we asked whether the prolonged activation seen in the frame-wide fluorescence signal was also visible in single cells, or whether individual neurons were active during different task events. Neuronal firing was sparse during the task, but we found that individual neurons were active during particular events in the task (Figure 4D,G). For instance, the red color-coded cell illustrated in Figure 4C and D became active soon after tone onset, whereas the neuron color-coded in gray fired during the post-reward licking period. The timing of their activities with respect to specific trial events seemed relatively stable, resembling what has been reported before for neurons in the striatum of behaving rodents by recording and analyzing spike activity (*Bakhurin et al., 2017; Barnes et al., 2011; Gage et al., 2010; Jog et al., 1999; Rueda-Orozco and Robbe, 2015*). To determine task encoding by single neurons at a population level, we defined task-modulated neurons as those that were active during any epoch of the task (see Materials and methods). Overall, 38.2% of the striatal neurons imaged in our samples were task-modulated. Of these, most were active during only one of the three task epochs (85%). Among task-modulated neurons, most were selectively active during the post-reward licking period (57%), but substantial numbers of neurons were also active during the tone presentation (17%) or after the licking had stopped (11%, Figure 4E,F).

Analysis of session-averaged population responses of neurons selectively active during the three epochs demonstrated a similar sequence of neuronal events as that which we found with analysis of the frame-wide fluorescence signals. The activation of a small group of neurons after cue onset was followed by a prolonged increase in responses of neurons active during the post-reward licking period (Figure 4F,G). This population activity ramped up until mice stopped licking, then quickly subsided (Figure 4F). Analysis of single-cell responses also identified a group of neurons that became maximally active following the end of licking. Grouping neurons based on the epoch during which they were active and sorting responses within each group by the timing of their peak session-averaged activity exposed a tiling of task time by neurons active in each of the three epochs (Figure 4G). Notably, the prolonged ramping of population activity observed during the post-reward licking period was produced by individual neurons being active within different specific time intervals during licking, not by their being active throughout the licking period.

### Striosomes encode reward-predicting tones more strongly than the matrix

To dissociate the specific contributions of striosomes and matrix to task encoding, we again first investigated aggregate neuronal responses in both striatal compartments. We drew region of interests (ROIs) around striosomes and around regions of the matrix with similar overall intensity of fluorescence and size, and then compared the total amount of fluorescence from these regions in trained animals. This analysis demonstrated a stronger tone-evoked activation in striosomes than in the nearby matrix regions sampled (Figure 5A) (ANOVA main effect p < 0.001). Moreover, the high-probability tone cue evoked a larger response than the low-probability tone cue (p < 0.001), and there was a trend for an interaction between compartment and tone (p = 0.055).

**Figure 5.**
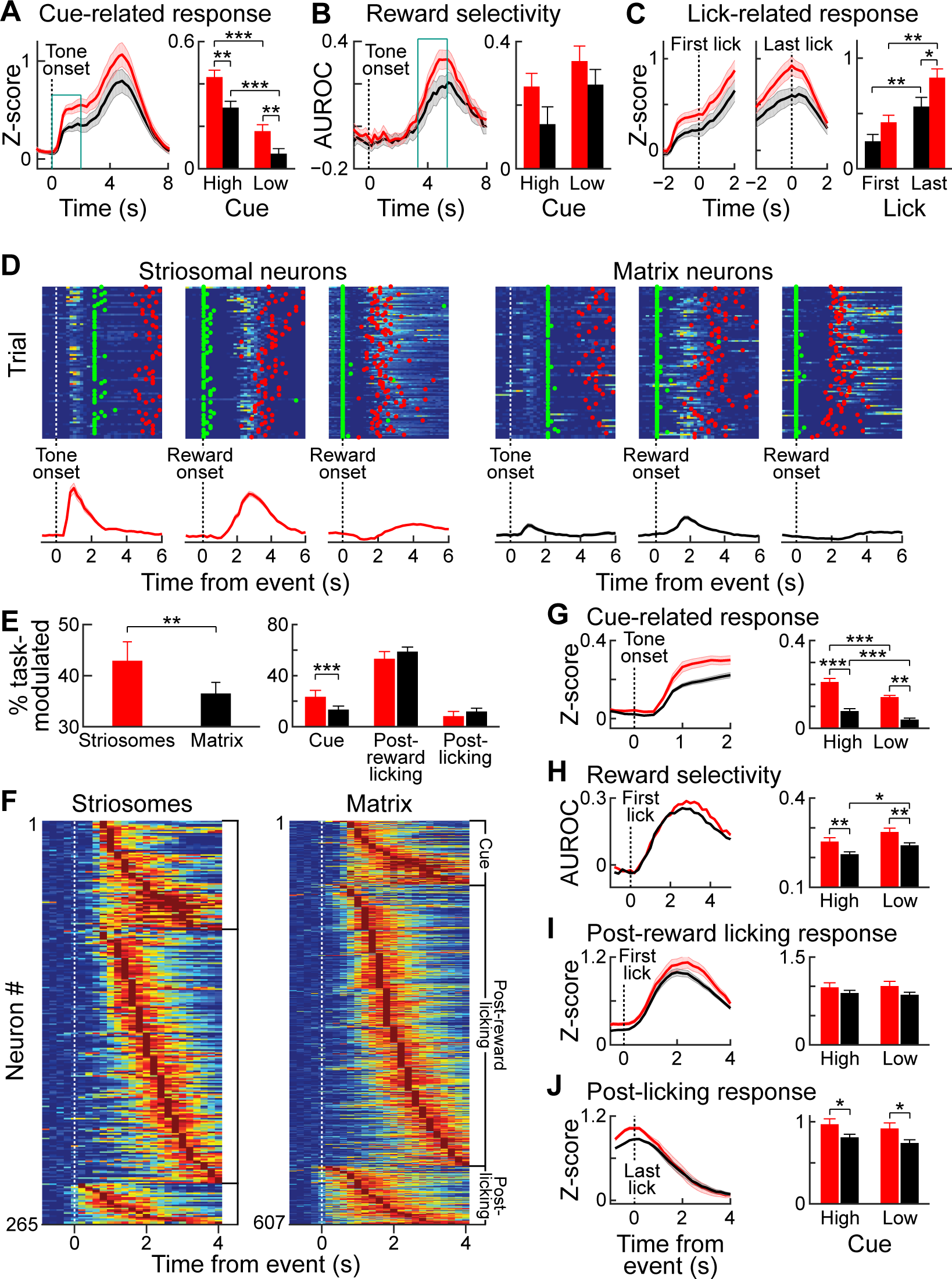
Striosomal cells respond more strongly to reward predicting cues than matrix cells. (**A**) Average striosomal (red) and matrix (black) neuropil activation during rewarded trials with high-probability cue (left), and quantification of the magnitude of the response to high‐ and low-probability cues (right), calculated for the time period indicated by blue box. **p < 0.01, ***p < 0.001 (ANOVA and post hoc t-test). Shading and error bars represent SEM. (**B**) Neuropil selectivity for the rewarded vs. unrewarded trials for every time point in high-probability trials (left) and the average selectivity during the time indicated in the blue box for both trial types (right). (**C**) Average neuropil post-reward activity aligned to the first (left) or last lick (middle), and average response during the ±1-s period. *p < 0.05, **p < 0.01 (ANOVA and post hoc t-test). (**D**) Trial-by-trial response of three striosomal (left block) and three matrix (right block) neurons that were selectively active during the cue (left), reward consumption (middle), or end of licking (right) task-epochs. Green and red dots show, respectively, the first lick after reward delivery and the last lick. Average responses for the same neurons are shown under the color plots. (**E**) Proportion of all task-modulated striosomal and matrix neurons (left) and those that were modulated selectively during cue, post-reward licking, or post-licking epochs of the task (right). **p < 0.01, ***p < 0.001 (Fisher's exact test). Error bars represent 95% confidence intervals. (**F**) Session-averaged responses of all task-modulated striosomal (left) and matrix (right) neurons. Neurons are grouped and sorted similar to those shown in Figure 4G. (**G**) Population-averaged responses of all task-modulated striosomal and matrix neurons to the high-probability cue (left), and the population responses separately averaged for high‐ and low-probability cues. **p < 0.01, ***p < 0.001 (ANOVA and post hoc t-test). Shading and error bars represent SEM. (**H**) Discriminability between rewarded and unrewarded trials for striosomal and matrix neurons. Left plot shows selectivity during trials with high-probability cue, and right plot shows average discriminability for all trials (quantified over 1-2 s time window after reward delivery). *p < 0.05, **p < 0.01 (ANOVA and post hoc t-test). (**I,J**) Population-averaged response during post-reward licking (**I**) or post-licking (**J**) periods, with data aligned to first and last lick after reward delivery, respectively. *p < 0.05 (ANOVA and post hoc t-test).

We tested for the selectivity of the responses to the high‐ and low-probability tone cues by quantifying the area under the receiver operating characteristic curve (AUROC) for these responses. Both striosomal and matrix responses displayed significant selectivity for the high-probability tone (p < 0.05), but there was no difference in selectivity between the striosomes and matrix at this stage of learning. We also performed an AUROC analysis to quantify the selectivity for rewarded trials (Figure 5B). Both striosomal and matrix neuropil had elevated activity in rewarded trials compared to non-rewarded trials during trials with both high‐ and low-probability tones. Repeated measures ANOVA showed that striosomes have a higher selectivity than the matrix for rewarded trials. Moreover, the selectivity for reward was larger in low-probability tone trials than high-probability tone trials. However, there was no interaction between cell-type and selectivity for reward (Figure 5B). Thus, both striosomes and matrix were more activated when the reward was less expected. We also investigated how the start and end of licking were reflected by activity in the two compartments (Figure 5C). The striosomal activation was higher than matrix activation (ANOVA main effect p < 0.001), and the activation was larger at the end of licking than in the beginning (Figure 5C) (±1 s around event, ANOVA main effect p < 0.001).

Next, we analyzed single-cell responses of striosomal and matrix neurons during the task. Individual neurons in both compartments were active during the task (Figure 5D). Overall, we found a higher proportion of striosomal neurons (42.9%; 312 out of 712 neurons) than matrix neurons (36.5%; 721 out of 1977 neurons) that were task-modulated (p < 0.005, Fisher's exact test; Figure 5E). Among the task-modulated neurons, a higher percentage of striosomal neurons were active during the cue epoch of the task (23.7% of striosome, 13.7% of matrix, p < 0.001, Fisher's exact test). By contrast, we found no differences in the percentages of striosomal and matrix neurons that were active during post-reward licking or post-licking (p > 0.05). In plots of session-averaged activity sorted by the timing of peak responses, we observed that striosomal and matrix neurons similarly tiled the temporal space of the task during each of the three epochs (Figure 5F). To compare responses of all task-modulated striosomal and matrix neurons during these epochs, we analyzed population-averaged activity aligned to different task events (Figure 5G-J). As in our neuropil analyses, we found that individual striosomal neurons were more robustly active than individual matrix neurons during the cue epoch of the task (Figure 5G, ANOVA main effect p < 0.001). Moreover, the high-probability tone elicited a higher response than the low-probability tone (p < 0.001).

We used an AUROC analysis to compare activity in trials that were rewarded (aligned to first lick after reward delivery) or unrewarded (aligned to 2 s after cue onset, a time period matching that for the rewarded trial analysis). We found that striosomal neurons were more selective for rewarded trials (Figure 5H, p < 0.001). Interestingly, the selectivity for reward was greater for low-probability than for high-probability tone trials (p < 0.01). While cells in the two compartments responded similarly during post-reward licking (Figure 5I, p > 0.05), striosomes had a higher response during the post-licking period (Figure 5J, p < 0.01). Together, these findings demonstrate that striosomes are more task-modulated in this appetitive classical conditioning task than the nearby matrix and that they are particularly more active during reward-predicting cues.

### Striosomal tone-evoked responses are acquired during learning

To determine how these responses were shaped by training, we analyzed striatal activity during the acquisition period of the task. To quantify levels of learning, we tested for significance in the difference between anticipatory licking for high‐ and low-probability cues during the tone presentation and the reward delay. If mice exhibited a significant difference on 2 out of 3 consecutive days, we considered them as being trained. Sessions performed before this criterion was met were categorized as acquisition sessions. This categorization allowed us to ask whether the strong striosomal cue-related response was a sensory feature, or whether it was an acquired response related to the meaning of the stimulus. Activity measures for the neuropil signals, comparing signals for all sessions before the mice reached the learning criterion with signals of all the sessions afterwards, showed that the response to the tones in striosomes was much stronger after animals learned the task (Figure 6A). The neuropil signal in striosomes was significantly higher after the task performance reached the training criterion (p < 0.05). This effect was not observed in the matrix (p > 0.05; ANOVA interaction p < 0.001). Single-cell analysis further indicated that during training, the percentage of task-modulated neurons increased steadily (Figure 6B), and that when mice reached the learning criterion (sessions 11-12 for the mice shown in Figure 6B,C), there was a rapid increase in the proportion of cue-modulated neurons (Figure 6C).

**Figure 6.**
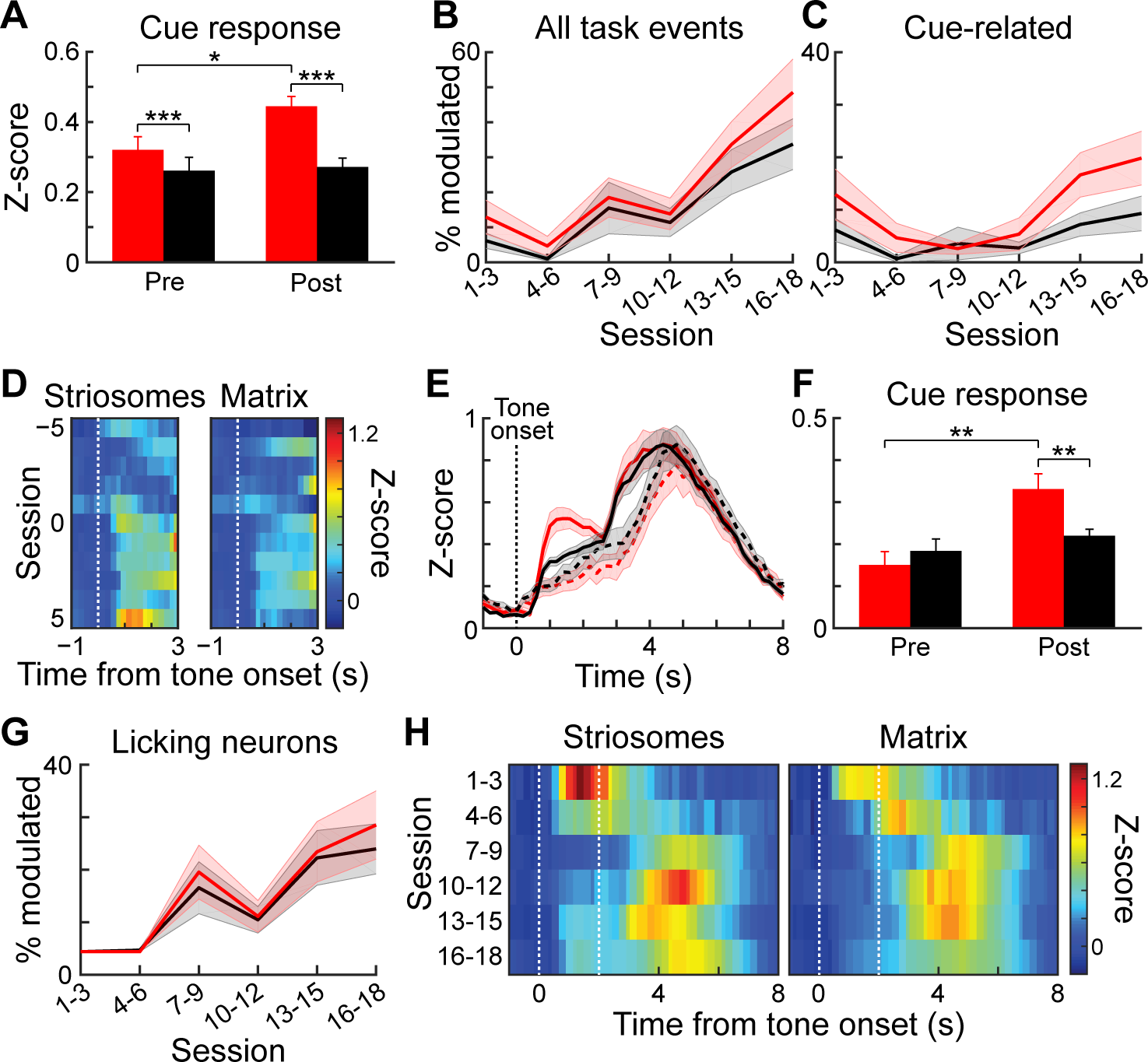
Cue-related signals in striosomes develop during training. (**A**) Average total striosomal (red) and matrix (black) neuropil signal during the 1.5-s tone period in all sessions before and after reaching learning criterion. *p < 0.05, ***p < 0.001 (ANOVA and post hoc t-test). Error bars represent SEM. (**B**) Percentage of task-modulated neurons in striosomes and matrix during the course of training. Shading represents SEM. (**C**) Percentage of cue-modulated neurons, shown as in B. (**D**) Mean z-score activity of task-modulated neurons during ±5 sessions around the session in which the learning criterion was reached (session 0). (**E**) Activity of task-modulated striosomal and matrix neurons averaged and normalized for blocks of 5 sessions before (dotted lines) and after (solid lines) reaching the learning criterion. (**F**) Quantification of the mean response of all task-modulated neurons during period from tone onset to reward onset. **p < 0.01 (ANOVA and post hoc t-test). (**G**) Percentage of licking-modulated neurons during training. (**H**) Mean normalized (z-score) activity of task-modulated neurons during training.

We further tested whether there was a sudden step-like increase in striosomal tone signaling. We averaged the z-scores of the activity of all task-modulated neurons during the last 5 sessions before and during the first 5 sessions after the learning criterion had been reached (Figure 6D). Averaging the activity in these two groups of sessions indicated a clear increase in striosomal signaling during the tone (Figure 6E). This increase was significant (Figure 6F,ANOVA, training main effect p < 0.001; interaction p < 0.05). In addition to this development of a tone response later in training, there was a tone-related activation in the sessions in which the animals were first exposed to the task, perhaps reflecting a startle or novelty signal effect. This tone-related activation disappeared after 1-3 sessions and reemerged later as mice learned the task. Analysis of neuronal activity during the post-reward licking period showed that during training there was an increase in the percentage of neurons that responded during this period (Figure 6G). Averaging the activity of all task-modulated neurons during training showed that there was an increase of activity in the period after reward delivery (Figure 6H). In contrast to the increases in tone response, this reward-period increase occurred several sessions before mice learned the task.

### During overtraining, striosomal tone-related responses intensify and become more selective for high-probability tones

To investigate further the relationship between neuronal responses and learning, two mice were trained for an additional five sessions. In these sessions, we imaged again the same fields of view that were recorded after these two mice reached training criterion. These last 5 sessions were considered overtraining sessions. The tone-evoked aggregate response became notably higher and sharper during this phase (Figure 7A). By contrast, the calcium signals that occurred immediately after reward delivery declined, resembling previously reported task-bracketing patterns (*Barnes et al., 2005; Jin and Costa, 2010; Jog et al., 1999; Smith and Graybiel, 2013; Thorn et al., 2010*). The increase in responses related to the tone during overtraining was particularly strong in striosomes. In the period following reward delivery, the signal initially dropped compared to the earlier sessions but subsequently reached the same magnitude. We quantified the peak response during the tone presentation period for the training, post-training and overtraining sessions (Figure 7B), comparing the responses of striosomal and matrix samples, and found a highly significant interaction (ANOVA interaction p < 0.005). In the trained and overtrained mice, striosomes had significantly higher tone-evoked responses than did the matrix (paired t-test, trained mice p < 0.01 and overtrained mice p < 0.05). The striosomal neuropil responses also became more selective for the high-probability cue during overtraining (Figure 7C), so that during overtraining the striosomal response was significantly larger than the matrix response (paired t-test p < 0.05).

**Figure 7.**
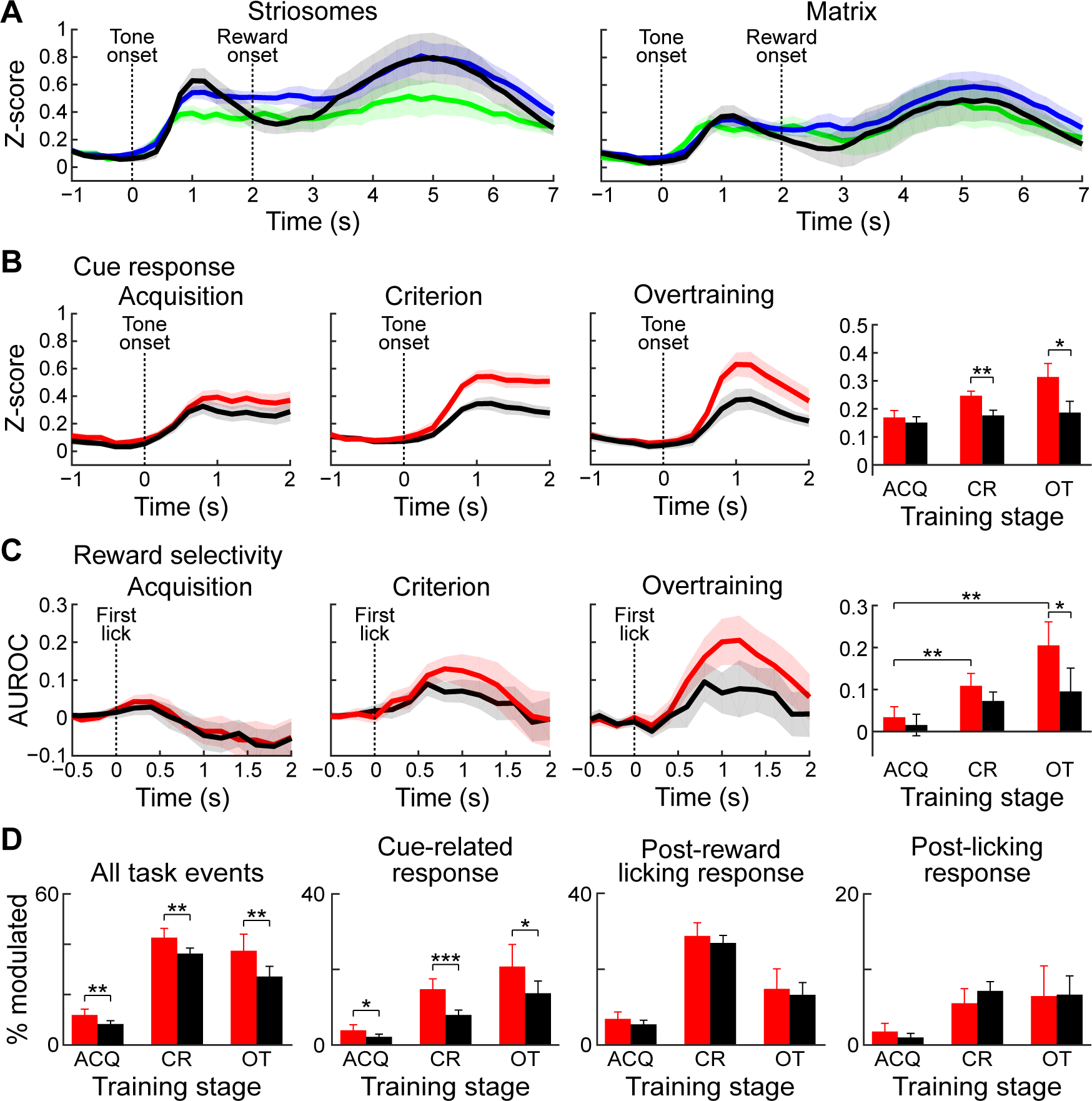
Striosomal cue-related responses strengthen during overtraining and become more selective. (**A**) Mean neuropil signals during training (green), after training (blue) and during overtraining (black) in striosomes (left) and matrix (right). Shading represents SEM. (**B**) Mean neuropil responses in striosomes and matrix during training (Acquisition), after training (Criterion) and during overtraining, and a quantification of the size of the response (right). *p < 0.05, **p < 0.01 (ANOVA). Shading and error bars represent SEM. (**C**) Selectivity for the high-probability cue, shown as in B. *p < 0.05, **p < 0.01 (ANOVA). (**D**) Percentages of task-modulated neurons (left), tone-modulated neurons (second), neurons modulated in post-reward licking period (third) and neurons modulated in post-licking period (right) during training (ACQ), after training (CR) and during overtraining (OT). *p < 0.05, **p < 0.01, ***p < 0.001 (Fisher's exact test).

We next analyzed the number of task-modulated neurons during the overtraining period. This percentage grew with training, then slightly dropped again during overtraining (Fig 7D, first panel; striosomes: 12.2% during training, 42.9% after training and 37.7% during overtraining; matrix: 8.6% during training, 36.6% after training and 27.5% during overtraining). At all stages, there were more striosomal than matrix task-modulated neurons (Fisher's exact test, p < 0.01). By contrast, the percentage of cue-modulated cells (Figure 7D, second panel) grew further (striosomes: 4.1% during training, 15.0% after training and 21.1% during overtraining; matrix: 2.3% during training, 8.1% after training and 13.9% during overtraining). There were more tone-modulated neurons during acquisition, after training and during overtraining (Fisher exact test, p < 0.05). The percentage of cells that were active during the consummatory licking period (Figure 7D, third panel) also increased (striosomes: 7.0% during training, 28.9% after training and 14.9% during overtraining; matrix: 5.6% during training, 27.0% after training and 13.3% during overtraining), but there was no difference between striosomes and matrix (Fisher's exact test, p > 0.05). The percentage of cells that were active after the end of licking remained stable during overtraining (Figure 7D, fourth panel; striosomes: 1.9% during training, 5.6% after training and 6.6% during overtraining; matrix: 1.1% during training, 7.3% after training and 6.8% during overtraining), with no differences between striosomes and matrix at any training stage (Fisher's exact test, p > 0.05). At no stage during training was the percentage of neurons that were significantly modulated during the response period or after the last lick different between striosomal and matrix neurons (Fisher exact test, p > 0.05). The limited number of significantly modulated neurons in these two mice was too low to make statistical comparisons of the neuronal responses. Nevertheless, the findings for the entire performance period of the mice collectively demonstrate that the activity patterns observed after training were acquired as mice learned the task, that the striosomal encoding of the tone became stronger than that of the matrix, that this response emerged at the time the animals began differentially responding to the tones, and that this response developed further during overtraining, becoming larger and more selective for the high-probability cue.

### Matrix responses are more sensitive to outcome history

In the classical conditioning task employed in this study, mice used the auditory tone presented during the cue epoch to guide their expectation for receiving a reward on the current trial. We examined their licking responses as a proxy for such expectation in order to ask whether, in addition to the information provided by the cue, the mice used the outcome in the previous trial to tailor their reward expectation in the current trial. In trials following rewarded trials, mice showed increased anticipatory licking during the cue and reward delay (Figure 8A; n = 33 sessions from five mice; p < 0.001, Wilcoxon signed-rank test), but licking during the post-reward period was unaffected by outcome on the previous trial (p > 0.05, Wilcoxon signed-rank test). To determine whether the task-related activity of the striatal neurons in our sample was also modulated by outcome history, we compared activity in trials preceded by a rewarded trial or by an unrewarded trial, regardless of the cue type (high or low probability) presented on the current trial. We first analyzed the effect of reward history on the cue-period responses of single task-modulated neurons and found that activity was slightly greater when the previous trial was rewarded (mean z-scores: 0.21 ± 0.01 vs. 0.17 ± 0.01 for previously rewarded and unrewarded; p < 0.01). When we analyzed the effect of outcome history on neural responses observed during post-reward licking in currently rewarded trials, we found that the activity of a subset of striatal neurons was highly sensitive to outcome in previous trials (Figure 8B). We observed enhanced activity during post-reward licking when the previous trial was unrewarded, compared to when the previous trial was rewarded. Similarly, population-averaged responses of task-modulated neurons were significantly higher when the previous trial was unrewarded, as compared to when it was rewarded (p < 0.001, Wilcoxon rank-sum test). Importantly, post-reward licking behavior was invariant to previous trial outcome, making it unlikely that the observed changes in neural activity were related to changes in the motor output during reward consumption.

**Figure 8.**
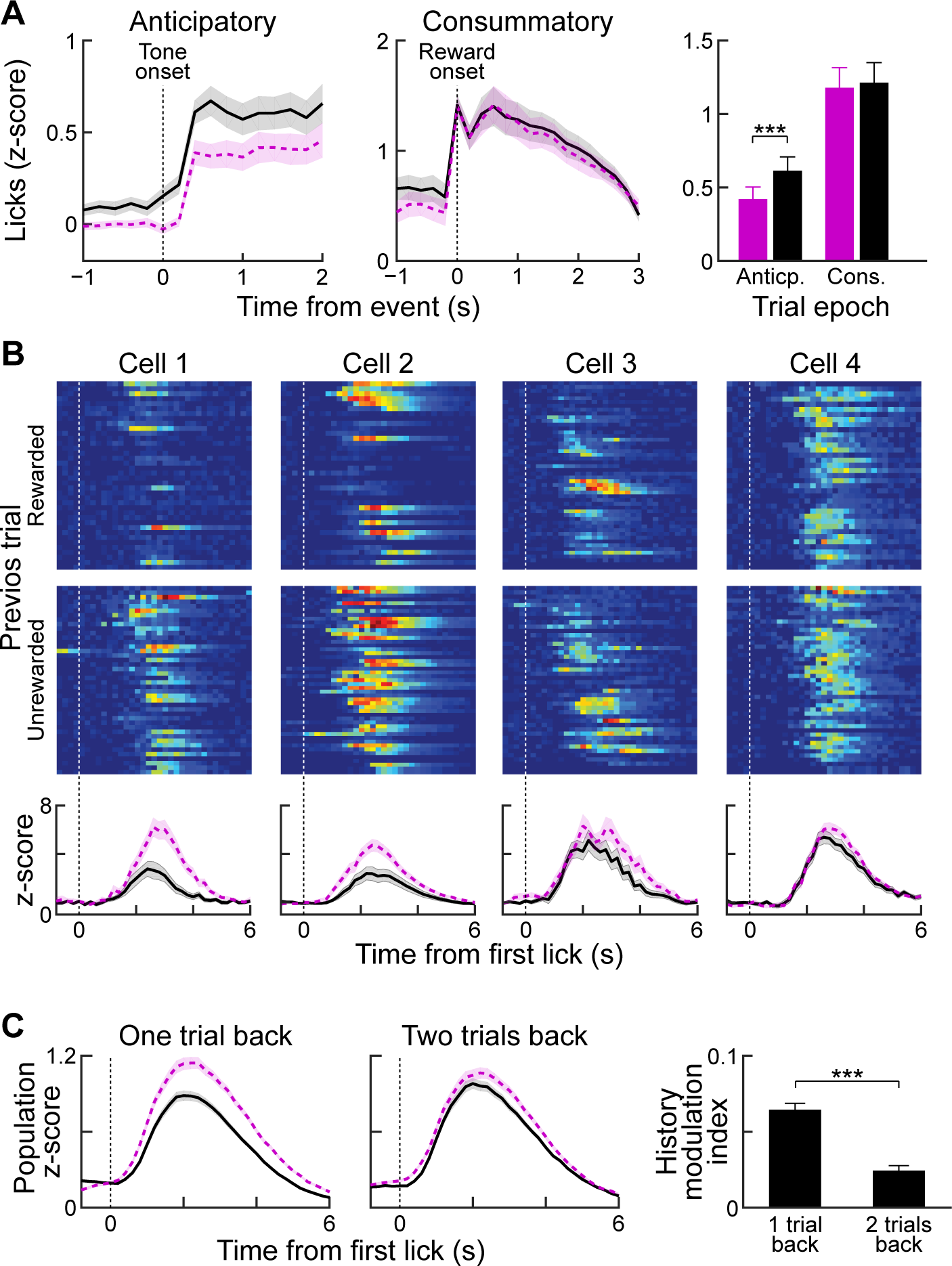
Reward history modulates anticipatory licking behavior and lick-period responses in striatal neurons. (**A**) Session-averaged licking activity during anticipatory and consummatory periods for trials in which the previous trial was rewarded (black solid) or unrewarded (purple dotted). Bar plot shows modulation of anticipatory or consummatory licking activity by reward history. ***p < 0.001 (Wilcoxon signed-rank test). Shading and error bars represent SEM. (**B**) Single-trial (top rows) and averaged (bottom row) post-reward licking responses of four sample neurons for previously rewarded (solid) or unrewarded (dotted) trials. (**C**) Population-averaged, post-reward licking activity of all task-modulated neurons for reward histories extending one or two trials back. ***p < 0.001 (Wilcoxon signed-rank test).

To determine how far back in time we could detect an outcome history effect, we computed a history modulation index (see Materials and methods) for currently rewarded trials with two types of reward history. In the first group, we separated rewarded trials based on whether the previous trial was rewarded or unrewarded (one trial back). For the second group, we disregarded the outcome status in the immediately preceding trial and separated trials depending on the outcome status of two trials in the past (two trials back). This analysis showed that recent reward history has a stronger influence on post-reward licking responses of task-modulated neurons than trials farther back in the past (Figure 8C, p < 0.001, Wilcoxon signed-rank test).

We asked whether this history modulation effect was detectable for both striosomal and matrix neurons (Figure 9). Examination of both population-averaged responses (Figure 9A) and single-cell responses (Figure 9B) suggested that both striosomal and matrix neurons were modulated by previous reward history, but that matrix neurons were more sensitive to this modulation. Quantification of this comparison by calculating the history modulation indices for striosomal and matrix neurons confirmed that the matrix responses were more influenced by previous reward history than the responses of striosomal neurons (Figure 9C,D; p < 0.01, Wilcoxon rank-sum test).

**Figure 9.**
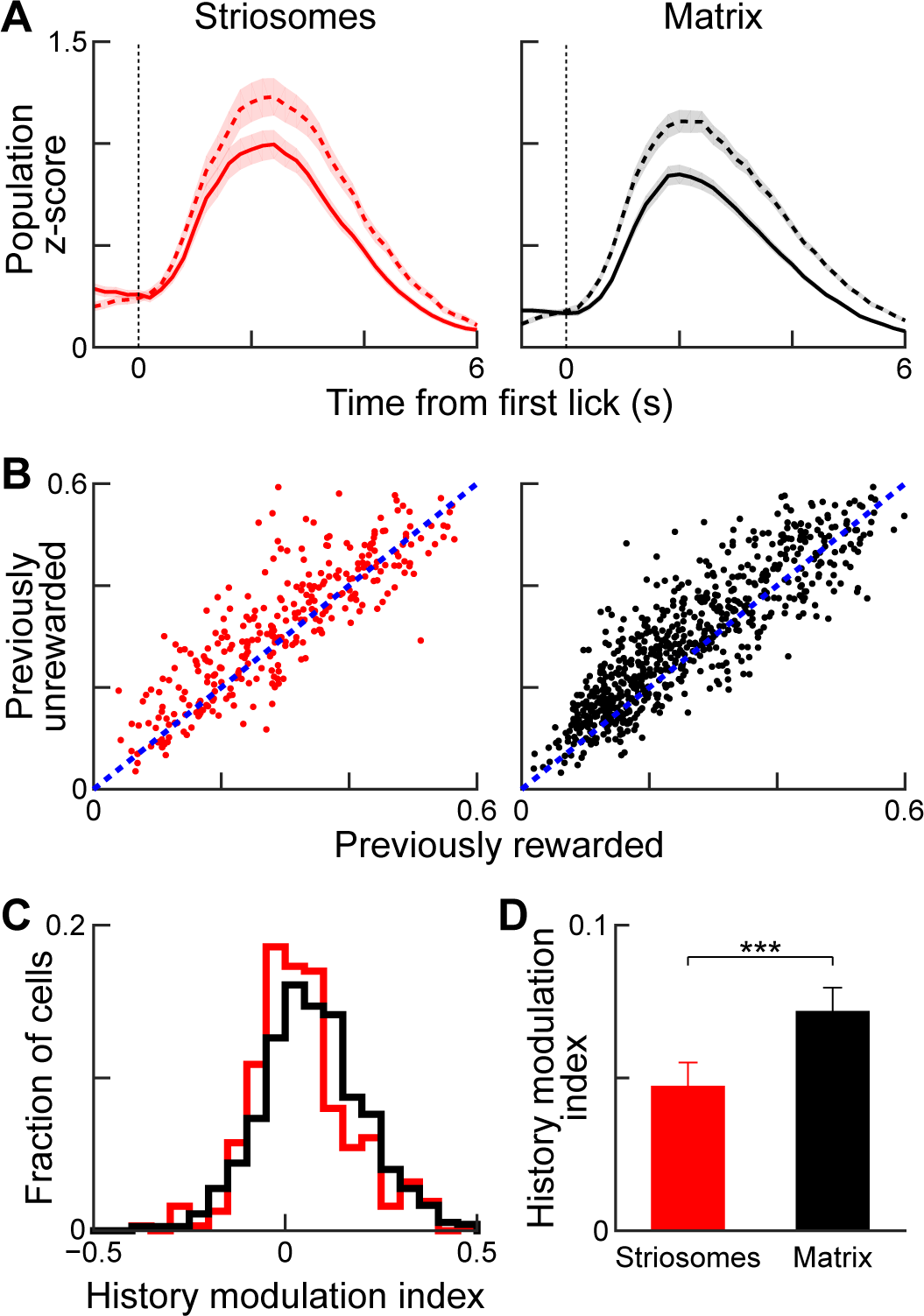
Reward-history modulation of striosomal and matrix neurons. (**A**) Population-averaged responses of all task-modulated striosomal (left) and matrix (right) neurons during trials following previously rewarded (solid) or unrewarded (dotted) trials. Shading represent SEM. (**B**) Normalized lick-period responses (averaged over 1-3 s after first lick) of individual striosomal (left) and matrix (right) neurons. Responses with previously rewarded trials are plotted against responses from previously unrewarded trials. Unity line is shown as blue dotted line (**C**) Histogram showing reward-history modulation index for all task-modulated striosomal and matrix neurons. (**D**) Mean reward-history modulation index for striosomal and matrix neurons. ***p < 0.001 (Wilcoxon rank-sum test). Error bars represent SEM.

## Discussion

Our findings demonstrate that 2-photon calcium imaging can be used to identify the activity patterns of subpopulations of striatal neurons distinguished as being in either the striosome or matrix compartments. Even with the use of a simple classical conditioning task involving cues predicting high or low probabilities of receiving reward, we could detect in all mice many task-related striatal neurons, altogether 38% of the 2704 neurons successfully imaged in the post-training phase. Among these, we found clear differences in the response of the striosomal and matrix neurons during cue presentation, found contrasts in the timing and selectivity of striosomal and matrix responses during learning and overtraining, and found that the responses of the two compartments were differentially affected by reward history. These findings, based on direct visual detection of striosomes by their birthdate-labeled neuropil and cell bodies, demonstrate that neurons of the two main compartments of the striatum have distinguishable response properties and likely encoding functions related to reinforcement learning and performance.

### Tone-period activity

By the time when the animals had reached the learning criterion, striosomes, examined both by averaged neuropil measures and by single-cell activities, were more responsive to the task than the surrounding matrix neurons. The differential activation of striosomes was particularly striking for the reward-predictive cues. More striosomal neurons were active in relation to the cues, and this effect grew stronger as animals learned. The striosomal neurons also were more selective for the high-probability cue. These responses did not reflect an overall greater response of striosomes to all conditions; for example, their responses were less sensitive than those of the matrix neurons to immediate reward history.

### Outcome period activity

Over the task-related population, the highest activity levels for many of the neurons as the learning criterion was reached occurred during the outcome period, whether the neurons were in striosomes or in the surrounding matrix. During this period, overall neuronal activity built up and peaked at the end of the licking. However, several factors pointed to this response as being different from a pure motor response to the licking movements. Most strikingly, even among the neurons strongly active during the prolonged licking period, the majority rose to their peak activity at specific times within this period rather than during the entire licking period. These post-reward peak responses tiled the entire time after reward. A subgroup of these neurons even peaked in activity after the end of the last licks. Second, we found dissociations between licking behavior and neuronal responses. For instance, activations during licking were larger when the previous trial was rewarded, whereas the licking behavior itself was not different. The anticipatory licking during the cue period and the neuronal responses during the cue also appeared dissociable during overtraining. During high-probability cues, when animals licked throughout the reward delay period, the neuronal signal decayed, whereas the opposite occurred during low-probability cues. Thus, although the signals observed during periods of licking are likely to be related to licking, their patterns of occurrence suggest an interesting multiplexing of information about licking, reward history, timing with respect to task events, and reward prediction. Importantly, we found the same stronger tone modulation in striosomes when we analyzed neuropil activity. In these analyses, we obtained matched, simultaneously registered striosomal and matrix data points from every session during the same behavioral performance. For this reason, the differences that we observed in the activity of the striatal compartments cannot be related to differences in licking behavior, as the behaviors were identical.

### Sensitivity to reward history

In contrast to these accentuated responses of striosomes, the striosomal neurons as a population were less sensitive than those in the matrix to immediate reward history. When the learning criterion had been reached, the neuronal responses for a given trial were elevated when the previous trial was not rewarded. By contrast, anticipatory licking was decreased in trials following unrewarded trials. These effects were significantly larger for the matrix. This reward history effect did not occur for two-back reward history, suggesting that it reflected immediate reward history.

### Learning-related differences in the responsiveness of striosomal and matrix neurons

Our recordings during the course of training demonstrated that both the cue-related responses and the post-reward responses were built up during behavioral acquisition of the task, with tone-related responses abruptly appearing when the mice reached the learning criterion. These learning-related dynamics suggest that the observed striosomal tone responses do not simply reflect auditory stimulus presentations. Moreover, during overtraining, the striosomal cue response strengthened: more striosomal neurons were significantly modulated by tone presentation, this striosomal response became stronger and more temporally precise, and it became more selective for the high-probability cue. By contrast, the activity in the period after reward delivery until the end of licking did not change notably and was even reduced slightly but non-significantly. Finally, the overall activity patterns in the neuropil began to resemble the classical task-bracketing pattern with peaks of activity at the beginning and the end of the trial (*Barnes et al., 2005; Jin and Costa, 2010; Jog et al., 1999; Smith and Graybiel, 2013; Thorn et al., 2010*).

In the matrix, the effects of overtraining were less pronounced. The response to the tone and reward consumption stayed similar but, as in the striosomes, a pattern resembling task-bracketing formed in the matrix. All of these effects could be detected not only at the single-cell level but also by assessing total fluorescence in defined striosomes and regions of the matrix with equivalent areas. These findings suggest that in striosomes, responses to reward-predicting cues are accentuated relative to responses detected in the matrix.

### Reward signaling in the dorsal striatum

It has previously been found that a minority of dorsal striatal neurons encode reward prediction errors (*Oyama et al., 2010; Oyama et al., 2015; Stalnaker et al., 2012*). Two of the major targets of striosomes, the dopamine-containing substantial nigra pars compacta and the lateral nucleus of the habenula, are well known to signal reward prediction errors (*Bayer and Glimcher, 2005; Bromberg-Martin and Hikosaka, 2011; Keiflin and Janak, 2015; Matsumoto and Hikosaka, 2007; Schultz, 2016; Schultz et al., 1997*). Therefore, we asked whether striosomes and matrix differentially encode reward prediction error signals. One particular possibility is that striosomes through their GABAergic innervation of dopamine-containing neurons could signal a negative reward prediction signal. However, we found that striosomes preferentially encoded reward-predictive cues. We did not find differences between striosomes and matrix in outcome-related activity. We also did not find signals related to reward omissions in either striosomes or matrix. We are aware that the dorsal striatum is heavily implicated in motor behavior, through learning, action selection or perhaps the invigoration of action (*Apicella et al., 1992; Balleine et al., 2007; Cui et al., 2013; Hikosaka et al., 2014; Kreitzer and Malenka, 2008; Mink, 1996; Nelson and Kreitzer, 2014; Packard and Knowlton, 2002; Redgrave et al., 1999; Samejima et al., 2005; Yin and Knowlton, 2006*). Nevertheless, we chose to start in these experiments by determining how a fundamental feature of the striatum, signaling of outcome and prediction of outcome, is represented in the striosome and matrix compartments. Future work will address the involvement of striosomes and matrix in action and decision-making.

### Striosome labeling

Visual identification of striosomes by their dense neuropil labeling was achieved by birthdating of striosomal neurons and their processes at the mid-point of striosome neurogenesis. Even though a minority of the striosomal neurons were pulse-labeled by the single tamoxifen injections, and despite the fact that there were scattered birthdate-labeled neurons in the extra-striosomal matrix, we could readily identify striosomes visually *in vivo* using 2-photon microscopy and in post-mortem MOR1-counterstained sections prepared to confirm the selectivity of labeling. We are aware that, with the use of pulse-labeling at neurogenic time points, we have incomplete labeling of compartments in any one animal, but the time of induction that we used was at the middle of the striosomal neurogenetic window and was days before the onset of major levels of matrix neuron neurogenesis (*Fishell and van der Kooy, 1987; Graybiel, 1984; Graybiel and Hickey, 1982; Kelly et al., 2017; Newman et al., 2015*). We are also aware that the matrix compartment itself is heterogeneous, but such heterogeneity could not be taken into account in our experiments. Further, our method did not rely on a single molecular or genetic marker to distinguish compartmental identify, but this had also a possible advantage in thereby avoiding potential unidentified biases that could arise from molecular-identity labeling. It is currently unknown to what extent there are different subtypes of striosomal neurons and what the exact neuronal subtype composition of striosomes is. Kelly et al. (*2017*) have found that at E11.5, both dopamine receptor D1 (D1R) and D2 (D2R) expressing neurons are being born, with a slight bias toward D1 neurons. However, other evidence suggests a predominance of D1R-containing neurons in striosomal mouse models (*Banghart et al., 2015; Cui et al., 2014; Smith et al., 2016*) or contrarily, a larger amount of D2R-containing neurons (*Salinas et al., 2016*). It is likely that differential labeling of subtypes of striosomal and matrix neurons occurs in different mouse lines, as has been seen by ourselves (J. Crittenden, personal communication, September 2017). It is clearly of great interest to determine neuronal response properties of specific subgroups of striosomal neurons as defined by genetic markers, but we here have chosen to have secure visual identification of striosomal and matrix populations based on the identification of restricted neuropil labeling of striosomes achieved by their birthdating and confirmed by their correspondence to the classic identification of striosomes in rodents as MOR1-dense zones (*Tajima and Fukuda, 2013*).

### Prospects for future work

Our findings are confined to the analysis of a very simple task, and they clearly are unlikely to have uncovered the range of functions of the striosome and matrix compartments. Yet the differences detected already suggest that striosomal neurons could be more responsive to the immediate contingencies of events than nearby matrix neurons, that they could gain this enhanced sensitivity by virtue of learning-related plasticity, but that they could be less sensitive to immediately prior reward history. These attributes of the striosomes could be related to real-time direction of action plans based on real-time estimates of value. To our best knowledge, this is the first report of simultaneous recording of visually identified striosome and matrix compartments in the striatum, here made possible by the neuropil labeling in pulse-labeled Mash1-CreER mice.

## Materials and methods

All experiments were conducted in accordance with the National Institute of Health guidelines and with the approval of the Committee on Animal Care at the Massachusetts Institute of Technology.

### Mice

Mash1(Ascl1)-CreER mice (*Kim et al., 2011*) (Ascl1tm1.1(Cre/ERT2)Jejo/J, Jackson Laboratory) were crossed with Ai14-tdTomato Cre-dependent mice (B6;129S6-Gt(ROSA)26Sor, Jackson Laboratory; (*Madisen et al., 2010*)) and crossed with FVB mice in the MIT colony to achieve tdTomato labeling driven by Mash1. Female Mash1-CreER x Ai14 mice were crossed with C57BL/6J males to breed the mice that we used for the experiments. Tamoxifen was administered by oral gavage (100 mg/kg, dissolved in corn oil) to induce Mash1-CreER at embryonic day (E) 11.5 in order to label predominantly striosomal neurons in anterior to mid-anteroposterior levels of the caudoputamen. Five mice (4 male and 1 female) were used for the imaging experiments.

### Surgery

#### Virus injections

Adult Mash1(Ascl1)-CreER x Ai14 mice received virus injections during aseptic stereotaxic surgery at 7–10 weeks of age. They were deeply anesthetized with 3% isoflurane, were then head-fixed in a stereotaxic frame, and were maintained on anesthesia with 1-2% isoflurane. Meloxicam (1 mg/kg) was subcutaneously administered, the surgical field was prepared and cleaned with betadine and 70% ethanol, and based on pre-determined coordinates, the skin was incised, the head was leveled to align bregma and lambda, two holes (ca. 0.5 mm diameter) were drilled in the skull. Two injections of AAV5-hSyn-GCaMP6s-wpre-sv40 (0. 5 μl each, University of Pennsylvania Vector Core) were made, one per skull opening, to favor widespread transfections of striatal neurons at the following coordinates relative to bregma: 1) 0.1 mm anterior, 1.9 mm lateral, 2.7 mm ventral and 2) 0.9 mm anterior, 1.7 mm lateral and 2.5 mm ventral. Injections were made over 10 min, and after a ~10-min delay, the injection needles were slowly retracted. The incision was sutured shut, the mice were kept warm with wet food during post-surgical recovery, and they were given meloxicam (1 mg/kg, subcutaneous) for 3 days to provide analgesia.

#### Cannula implantation

We assembled chronic cannula windows by adhering a 2.7-mm glass coverslip to the end of a stainless steel metal tubing (1.6-1.8 mm long, 2.7 mm diameter; Small Parts) using UV curable glue (Norland). Cannula windows were kept in 70% ethanol until used for surgery. At 20-40 days after virus injection, mice were water restricted, and a second surgery was performed under deep isoflurane anesthesia as before to allow insertion of a cannula for imaging (*Dombeck et al., 2010; Howe and Dombeck, 2016; Lovett-Barron et al., 2014*) and mounting of a headplate to the skull for later head fixation. Bregma and lambda were aligned in the horizontal plane, and the anterior and lateral coordinates for the craniotomy were marked (0.6 mm anterior and 2.1 mm lateral to bregma). The skull was then tilted and rolled by 5° to make the skull surface horizontal at the location of cannula implantation. A 3 mm diameter craniotomy was made with a trephine dental drill. The exposed cortical tissue overlying the striatum was aspirated using gentle suction and constant perfusion with cooled, autoclaved 0.01 M phosphate buffered saline (PBS), and part of the underlying white matter was removed. A thin layer of Kwiksil (WPI) was applied, and the chronic cannula was inserted into the cavity. Finally, metabond (Parkell) was used to secure the implant in place and to attach a headplate to the skull. The mice received the same post-surgical care as described above.

### Behavioral training

When mice had recovered from surgery and the optical window had cleared, they were water deprived (1-1.5 ml per day) and habituated to head-fixation for on average 5 days. During head fixation, the mice were held in a polyethylene tube that was suspended by springs. When they showed no clear signs of stress and readily drank water while being head-fixed, behavioral training was begun. Water was delivered through a tube controlled by a solenoid valve located outside of the imaging setup, and licking at the spout was detected by a conductance-based method (*Slotnick, 2009*). In the behavioral training protocol, 2 tones (4 or 11 kHz, 1.5-s duration) were played in a random order. The tones predicted reward delivery (5 μl) with, respectively, an 80% or 20% probability. In each trial, there was a 500-ms delay after tone offset before reward delivery. Inter-trial intervals were randomly drawn from a flat distribution between 5.25 and 8.75 s. Training was considered to be complete when there was a significant difference in anticipatory licking during the cue period (two-sided t-test, α= 0.05). Imaging was performed daily during training and continued for 3-7 sessions afterwards. Two mice were then given 5 overtraining sessions.

### Imaging

Imaging of GCaMP6s and tdTomato fluorescence was performed with a commercial Prairie Ultima IV 2-photon microscopy system equipped with a resonant galvo scanning module and a LUMPlanFL, 40x, 0.8 NA immersion objective lens (Olympus). For fluorescent excitation, we used a titanium-sapphire laser (Mai-Tai eHP, Newport) with dispersion compensation (Deep See, Newport). Emitted green and red fluorescence was split using a dichroic mirror (Semrock) and directed to GaAsP photomultiplier tubes (Hamamatsu). Individual fields of view were imaged using either galvo-resonant or galvo-galvo scanning, with acquisition framerates between 5 and 20 Hz. Laser power at the sample ranged from 11 to 42 mW, depending on GCaMP6s expression levels. For final analysis of the dataset, all imaging sessions were resampled at a framerate of 5 Hz.

Fields of view were chosen on the basis of clear labeling of putative striosomes defined by dense tdTomato signal in the neuropil. Within these zones, both tdTomato-positive as well as unlabeled cells were present and were defined as putative striosomal cells. Because of the 2.4-mm inner diameter of the cannula, we could typically find several striosomes that we could image at different depths. Our sampling strategy was to image as many different neurons as possible. During training, we rotated through the field of views, but after training and during overtraining we imaged unique, non-overlapping field of views.

### Image processing and cell-type identification

Calcium imaging data were acquired using PrairieView acquisition software and were saved into multipage TIF files. Data were analyzed by using custom scripts written in ImageJ (National Institutes of Health) or Matlab (Mathworks). Images were first corrected for motion in the X-Y axis by registering all images to a reference frame. We used the pixel-wise mean of all frames in the red channel containing the structural tdTomato signal to make a reference image. All red channel frames were re-aligned to the reference image by the use of 2-dimensional normalized cross-correlation (template matching and slice alignment plugin (*Tseng et al., 2011*)). The green channel frames containing the GCaMP6s signal were then realigned using the same translation coordinates with the ‘Translate’ function in ImageJ. After realignment, ROIs were manually drawn over neuronal cell bodies using standard deviation and mean projections of the movies. With custom MATLAB scripts, we drew rings around the cell body ROIs (excluding other ROIs) to estimate the contribution of the background neuropil signal to the observed cellular signal. Fluorescence signal for each cell was computed by taking the pixel-wise mean of the somatic ROIs and subtracting 0.7x the fluorescence of the surrounding neuropil, as previously described (*Chen et al., 2013*). After this step, the baseline fluorescence for each cell (F_0_) was calculated using K-means (KS)-density clustering to find the mode of the fluorescence distribution. The ratio between the change in fluorescence and the baseline was calculated as DFF = F_t_ – F_0_/F_0_.

Individual cells were identified as striosomal if their cell body was in a region that was densely labeled by tdTomato, or if the cells themselves were tdTomato-positive. Hence, the small minority of tdTomato neurons that appeared in the matrix (*Kelly et al., 2017*) was included in the striosomal population. Altogether 6320 neurons were recorded (during training: 2871; after training: 2704; overtraining: 745). Of these, 1867 were considered striosomal (during training: 912; after training; 727; during overtraining: 228). Of these, 294 were labeled with tdTomato, 1828 were located in densely tdTomato-labeled striosomes and 255 met both criteria. There were 39 tdTomato-labeled cells that were not located in a zone of dense tdTomato neuropil labeling. We excluded these neurons in multiple analyses, but their exclusion never resulted in a different outcome in our analyses.

### Analysis of neuronal activity

#### Neuropil analysis

To provide a first insight into striosomal and matrix signaling, we integrated the fluorescence signal from within an identified striosome and from a part of the matrix in the same field of view that had a similar size, background fluorescence and number of neurons. DFF, calculated as DFF = F_t_– F_0_/F_0_, was normalized by calculating z-scores relative to the signal at the end (1 s) of inter-trial intervals to correct for relative differences between sessions. To determine the selectivity of responses to different task events, the AUROC curves were calculated. For cue selectivity, we calculated the AUROC curve by comparing the response during high‐ and low-probability cues. For the selectivity to rewarded trials, we calculated the AUROC by comparing rewarded and unrewarded trials for the two cues separately.

#### Single cell analysis

The conditioning task had three epochs — cue, post-reward licking, and post-licking. To identify task-modulated neurons active during these epochs, we aligned the data either to tone onset, to the first lick after reward delivery, or to the end of licking. We compared the fluorescence values over the following time windows to a 1-s baseline preceding each event. For the tone-aligned data, mean fluorescence was calculated over a 2-s time window after tone onset separately for trials with either the high‐ or low-probability cues. Neurons that were significantly active in either of the cue conditions were considered to be task-modulated. To find neurons modulated during post-reward licking, GCaMP6 fluorescence was averaged between the time when the animal first licked to the reward and the time that it stopped licking. We also used a 1-s time window after end of licking for identifying task-modulated neurons during this period. In some trials, animals did not stop licking until start of the next trial. These trials were excluded from the analysis due to the difficulty in assigning licking end-time. For a neuron to be considered as task-modulated, we required that its activity exhibit a significant increase from baseline for any of the three alignments (two-sided Wilcoxon rank-sum test; α = 0.01, corrected for multiple comparisons). Neurons exclusively active during only one epoch of the task were considered to be selectively responsive during that period. Most neurons (>80%) were significantly active only during one of the epochs. To compare signals across neurons, we used z-score normalization of the DFF signals with a 1-s period before the cue as a baseline. For analysis of the peak activity of task-modulated neurons, DFF signals were normalized to the maximum of the session-averaged activity for any particular alignment in order to compare peak activity times during the time interval of interest.

To determine whether reward outcome in the previous trial modulated licking behavior during the task, we first compared anticipatory licking in trials that were followed by either rewarded or unrewarded trials. We included all current trials, regardless of the cue or the outcome status. To examine the effect of outcome history on licking after reward delivery, we analyzed only currently rewarded trials, again ignoring the identity of the cue presented. To determine whether neural responses were modulated by previous outcome history, we computed a history modulation index (HMI) using the following formula:

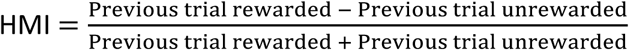

HMI was computed from z-scored values normalized by the following method. First, we took all currently rewarded trials and averaged the z-scored DFF values over a 2-s window period starting 1 s after reward delivery. We choose this time window because we found that most of the task-modulated neurons were active during this period. These values were then scaled by the range of observed responses, so that normalized values ranged from 0 to 1. Trials were then separated based on different outcome histories.

#### Statistical analysis

We used Wilcoxon sign-rank tests to test for significance modulation of single neurons in different task epoch. ANOVA was used for evaluating interactions between multiple factors. For percentages, Fisher's exact test was used to compare groups and confidence intervals were calculated using binomial tests.

### Histology

After the experiments, mice were transcardially perfused with 0.9% saline solution followed by 4% paraformaldehyde in 0.1 M NaKPO_4_ buffer (PFA). The brains were removed, stored overnight in PFA solution at 4oC and transferred to glycerol solution (25% glycerol in tris buffered saline (TBS) until being frozen in dry ice and cut in transverse sections at 30 μm on a sliding microtome (American Optical Corporation). For staining, sections were first rinsed 3 x 5 min in PBS-Tx (0.01 M PBS + 0.2% Triton X-100), then were incubated in blocking buffer (Perkin Elmer TSA Kit) for 20 min followed by incubation with primary antibodies for GFP (Polyclonal, chicken, Abcam ab13970, 1:2000) and MOR1 (Polyclonal, goat, Santa Cruz sc-7488, 1:500). After two nights of incubation at 4oC, the sections were rinsed in PBS-Tx (3 x 5 min), incubated in secondary antibodies Alexa Fluor 488 (donkey anti-chicken (Invitrogen), 1:300) and Alexa Fluor 647 (Alexa Fluor 647, donkey anti-goat (Invitrogen), 1:300) for 2 hr at room temperature, rinsed in 0.1 M PB (3 x 5 min), mounted and covered with a coverslip with ProLong Gold mounting medium with DAPI (Thermo Fisher Scientific).

## Acknowledgements

We thank Dr. Mark Howe and Dr. Dan Dombeck for invaluable advice on 2-photon imaging of the striatum; we thank Dr. Leif Gibb and Jannifer Lee for initiating the breeding program and Dr. Josh Huang and Dr. Sean for their advice in this process, and Cody Carter for critical help with the breeding of the mice; and we thank Erik Nelson for his work on histology.

## Additional information

Funding

**uTable 1.**
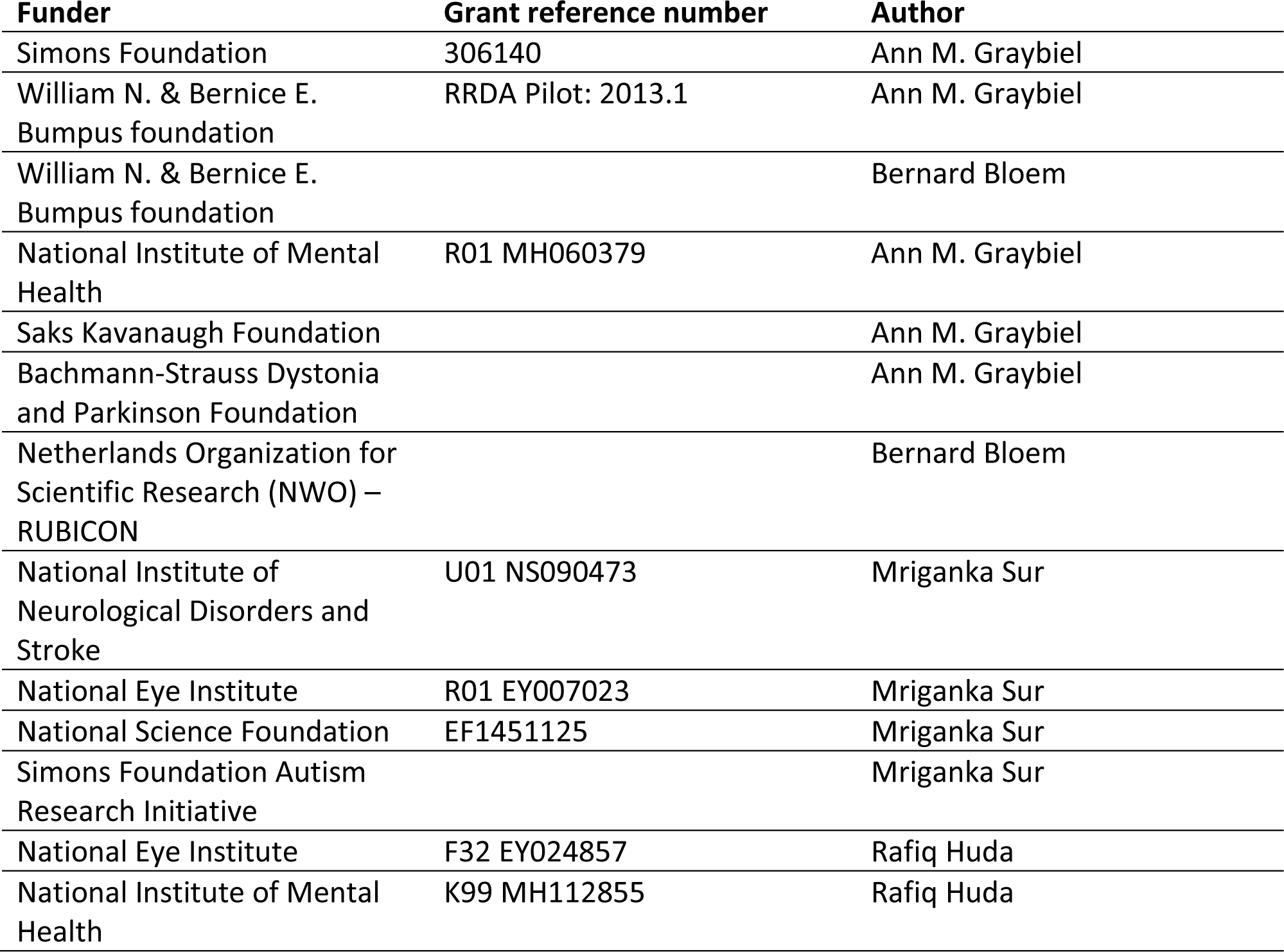
Funding.

## Author contributions

All authors participated in the design of the experiments; BB and RH performed imaging experiments; BB and RH performed analysis with input from AMG and MS; BB, RH and AMG wrote the manuscript with input from MS.

